# Nitrergic neurons of the forepaw representation in the somatosensory and motor cortices

**DOI:** 10.1101/2020.07.05.186635

**Authors:** Bárbara de Paula P. F. Guimaraes, Marco R. Curado, Anaelli Aparecida Nogueira-Campos, Jean Christophe Houzel, Ricardo Gattass

## Abstract

Nitrergic neurons (NN) are inhibitory neurons capable of releasing nitric oxide (NO) that are labeled with NADPHd histochemistry, allowing the study of their distribution and morphology. The rat primary somatosensory (S1) and motor (M1) cortices are a favorable model to investigate the morphology of the NN population. The distribution of the type I NN of the forepaw representation in the primary somatosensory (S1) and motor (M1) cortices of the rat in different laminar compartments and the morphological parameters related to the cell body and dendritic arborization were measured and compared. We found that the neuronal density in the S1 (130 NN/mm3) was higher than in the M1 (119 NN/mm3). Most NN neurons are multipolar (S1 with 58%; M1 with 69%) and a minority are horizontal (S1 with 6%; M1 with 12%). NN found in the S1 had a higher verticality index than those of the M1, and no statistical differences was found for the others morphological parameters. We also demonstrated statistical differences for most of the morphological parameters of the NN between different cortical compartments of the S1 and M1. Our results indicate that the NN of the forepaw in the S1 and M1 correspond to a single neuronal population whose functionality is independent of the different types of sensory and motor processing. However, the morphological differences found between cortical compartments of S1 and M1, as well as the higher density of NN found in the S1 indicate that the release of NO varies along and between the areas.

## 1. Introduction

Nitric oxide (NO), the first gaseous neurotransmitter described in the nervous system, is involved in many different physiological and pathological functions (Bredt & Snyder, 1992; Calabrese et al., 2007; Freire et al., 2012; Garthwaite, 2008; Moncada, 1991). In the brain, the nitric oxide synthases (NOS) enzymes that are capable of synthesizing NO are colocalized with nicotinamide adenine dinucleotide phosphate diaphorase (NADPHd) (Dawson et al., 1991; Stuehr, 2004). NADPHd histochemistry reveals the presence of two types of NO synthesizing neurons in the neuropil, type I and type II, characterized by a background composed of neurites and very thin reactive terminals (Franca et al., 2000). Type I neurons are intensely labeled, comprise 2% of cortical neurons and have been identified in all species studied until now. Compared to type I, type II neurons are weakly labeled, smaller and more numerous in the cortex with dendritic trees poorly or not labeled (Franca et al., 2000; Freire et al., 2004; Luth et al., 1994; Yan., 1997)

Using a neuronal reconstruction system (Nogueira-Campos et al., 2012) it is possible to investigate the distribution and morphology of these type I inhibitory neurons, known as nitrergic neurons (NN) in order to understand the microstructural characteristics of these neurons and their functionality between different cortical areas and layers, as observed with pyramidal neurons (Elston, 2002; Elston et al., 2006). Although some studies have investigated the nitrergic circuitry in the S1, M1, visual and auditory cortex, the hippocampus, the cerebellum, the striatum and the spinal cord (Franca et al., 2000; Freire et al., 2012; Garbossa et al., 2005; Nogueira-Campos et al., 2012; Santiago et al., 2007; Torres et al., 2006; Vlasenko et al., 2008), it is not well understood how this circuity structure is organized across the brain (Freire et al., 2012).

The mammalian primary somatosensory cortex (S1) and the primary motor cortex (M1) consist in two different cortical regions with interconnected sub-regions that have different functionalities thought to play distinct roles involving from sensory input to motor output, with the execution of voluntary movements (Morandell & Huber, 2017; Tandon et al., 2007). In the representation of the rat forepaw these areas share the same border, a useful and widely used model for exploring details of cortical organization in mammalian brains (Donoghue & Wise, 1982; Hall & Lindholm, 1974; Petersen, 2007). The S1 has six different layers from the pia mater to the white matter, with different laminar arrangements, and complete body representation (Rocha et al., 2007; Santiago et al., 2007). The M1 also has a distinct laminar arrangement in five different parallel layers from the pia mater to the white matter (Donoghue & Wise, 1982) and like the S1, has an organized topologically representation map (Donoghue et al., 1990; Hall & Lindholm, 1974; Tandon et al., 2007).

Investigating the cortical microstructure of the NN across their distinct cortical representations by comparing the distribution and fine morphological arrangement displayed by the cell bodies and dendritic arbors of the NN according to their position in the cortical circuitry can contribute to better understanding the structural and functional sensorimotor organization of this inhibitory nitrergic circuitry.

In the present study, we quantify and compare the distribution and morphology of the NN population in different laminar compartments of the forepaw representation in rat S1 and M1.

## 2. Materials and Methods

### Animals, perfusion, and tissue preparation

All experimental protocols were conducted following the NIH Guide for Care and Use of Laboratory Animals (www.nap.edu/catalog/12910.html) and were approved by the Ethics Committee for Animal Use in Scientific Research (CEUA) of the Center for Health Sciences (CCS) of the Federal University of Rio de Janeiro (UFRJ, protocol # IBCCF 029). Animals were obtained from the animal facilities of the Institute of Biological Sciences (ICB, UFRJ). All efforts were made to minimize suffering and to reduce the number of animals used.

Two adults male Wistar rats were deeply anesthetized and perfused through the left ventricle, using a Milan peristaltic pump, with 250 ml of 0.9% sodium chloride (NaCl), followed by 300–400 ml of 4% paraformaldehyde in pH 7.4, 0.1 M sodium phosphate buffer (PB). On the same day as the perfusion, the brain of each animal was removed from the skull, blocked, sectioned, and reacted for NADPHd histochemistry (see below). A block of the left hemisphere containing the entire frontal and parietal cortices was prepared and cut in the coronal plane with a vibratome (Leica model VT 1000S) into serial 150 μm-thick sections. The sections were collected in 0.1 M tris buffer, pH=8.0, and then reacted for NADPHd histochemistry.

### NADPHd histochemistry

The sections were incubated free-floating at 37°C in a solution consisting of 0.6% malic acid, 0.03% nitroblue tetrazolium, 1% dimethylsulfoxide, 0.03% manganese chloride, and 0.1% β-NADP in 0.1 M tris buffer, pH 8.0 (modified from Scherer-Singler et al., 1983). To increase penetration of reagents into the section thickness, the detergent Triton X-100 was added to the histochemical solution at a concentration of 0.75%. The sections were incubated in a shaker for 2-4 hours protected from light. The histochemical reaction was periodically monitored by inspecting one or two sections under the light microscope. When most of the NADPHd-reactive cell bodies in the section were presenting well-labeled tertiary dendrites, the histochemical reaction was stopped with tris buffer washes. Sections were then mounted onto poly-L-lysine-coated glass slides and left to air-dry overnight. The slides were then dehydrated in alcohol, washed twice in xylene for 5 min and coverslipped with Entellan (Merck).

### Delimitation of the S1, M1 and cortical compartments

The regions of interest in this study comprised the representation of the forepaw in the S1 and M1 (Figure 1a). According to Paxinos and Watson (1987), the representation of the forepaw in the S1 shares a common border with the M1 laterally for an anteroposterior extent of at least 1.0 mm. The portion of the M1 that borders the S1 contains the motor representation of the forepaw (Tandon et al., 2007) were defined with the aid of the adult rat brain atlas. Anatomical landmarks, such as the caudate-putamen, the anterior commissure, and the form and thickness of the corpus callosum, additionally helped to confirm the estimation of the anteroposterior level of the coronal sections used in this study. Because the border separating the M1 from the S1 is not clearly delineated absolutely sharp, a safety region of 200 µm in the mediolateral extent was adopted in the transition between these areas. To avoid mistakes in the attribution of the location of the NN cell body (in the S1 or M1) NN with cell bodies in the transition zone were not reconstructed, and thus not included in the analysis performed in this study.

**FIGURE 1.**
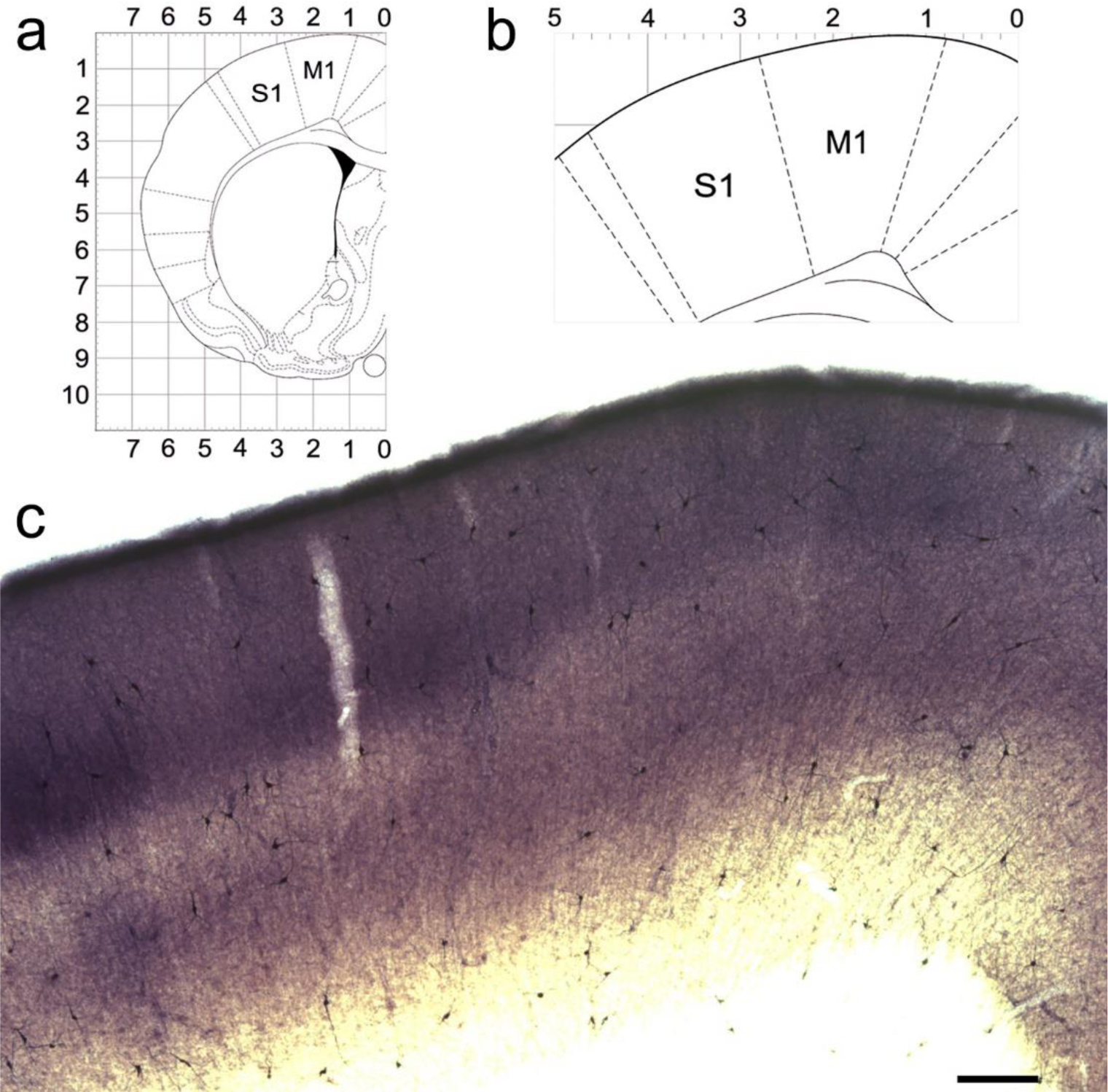
NADPHd reactive neuropil and nitrergic neurons. (a) Coronal section of the rat adult brain (Paxinos and Watson, 1998) at the level of the representation of the forepaw, and enlarged in (b). (c) Photomicrograph at the level of the representation of the forepaw showing NADPHd reactive neuropil and NN in areas S1 and M1. The representation of the forepaw in areas S1 and M1 are parallel to each other. In S1, the layer IV is more intensely labeled than layers I-III and V-VI. Scale bar: 200 μm.

Cortical areas and their laminar subdivisions were delimited using a 10x objective. The combination of a cortical area with a given cortical layer defined a cortical compartment in which we analyzed different quantitative parameters such as neuronal density, and different morphological characteristics such as cell body area, and dendritic length, among others (see below). Neuropil reactivity thus allowed the definition of the following laminar compartments in the S1 and the M1: the upper (supragranular) layers (UL, comprising layers I, II and III); layer V (V); layer VI (VI); and the lower (infragranular) layers (LL, comprising both layers V and VI together). The granular layer (GL or layer IV) was identified as an additional compartment exclusively in the S1. In order to calculate NN density, the limits of each of these compartments were drawn using Neurolucida 8.0 and Neuroexplorer software (MBFBiosciences, Inc), and these limits were then used to measure the compartment areas.

### Stereological evaluation of nitrergic neuronal density

Nitrergic neuronal density was evaluated in the representation of the forepaw in the S1 and M1 in the left hemisphere in the two animals that were used for neuronal morphological analysis. In each animal, we used a sequence of three adjacent 150-µm-thick sections reacted for NADPHd histochemistry, comprising a total thickness of 450 µm in each hemisphere. Neuronal density measurements in the different compartments of the S1 and M1 followed the procedures previously described by Nogueira-Campos et al (2012). Briefly, we counted the total number of nitrergic cell body profiles (*N*) and divided it by the flat area (*A*) of the cortical compartment. The compartment area was measured using the Neurolucida software to draw a quadrangle comprising the studied compartment, and then measuring the area of the quadrangle with Neuroexplorer. Density per compartment area was calculated after adding up all nitrergic cell body profiles found in a given compartment in all six sections used in this study; and then dividing this number by the sum of the compartment areas measured in each of those six sections.

Density per volume was then estimated for each cortical compartment adopting Abercrombie’s stereological correction formula (Abercrombie, 1946):

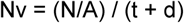

where *t* was the section thickness, and d was the average cell body diameter in each compartment. This diameter was estimated following the procedures described by Schuz and Palm (1989), assuming that t corresponded to the thickness set at the vibratome (150 µm). The diameter (d) was estimated from the cell body area (*a*), using the formula of the circle:

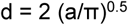

The average cell body diameter (*d*) was calculated for each cortical compartment using all reconstructed neurons located in that given compartment. Then, the value of *d* was corrected using the stereological estimations defined by formulae (2) and (3) of Schuz and Palm (1989); yielding values *d2* and *d3*, respectively. Since *d2* overestimates, and *d3* underestimates the actual value of *d* (Schüz & Palm, 1989), the final value of d applied in Abercrombie’s formula was the mean obtained from *d2* and *d3*.

### Tridimensional reconstruction

All NN identified in the regions under study were reconstructed, even when their dendrites spread out of the delimited areas. Neurons that presented evidence of being incomplete due to tissue sectioning were identified by the location of the cell body (usually close to the edge of the section, as estimated by the plane of focus) and by the presence of dendrites that abruptly disappeared at the edge of the section. Although these incomplete neurons were reconstructed for the quantification of neuronal density for illustration purposes, their morphometric data were not included in the quantitative analysis. Only complete neurons, with cell bodies located in the middle of the sections, with the most part of the cell bodies located within the delimited cortical compartments and with all dendrites presenting a tapering profile were included in the quantitative analysis of morphological parameters.

Digital reconstruction of NN was performed using both a Zeiss 40x neofluar plan objective (for the cell body) and a 100x plan objective (for the dendritic tree) coupled to a Zeiss Axioplan-2 microscope equipped with a color digital camera (1600×1200, 3/4” chip, 36bit, MBF), a motorized stage (Mac5000 LUDL), and an extra *z* encoder controlled by Neurolucida 8.0 software (MBFBiosciences, Inc) running on a Dell workstation.

### Acquisition of morphological parameters

A number of quantitative morphological parameters collected during neuronal reconstructions using the Neurolucida system were afterward extracted using NeuroExplorer software (MBFBiosciences, Inc), as previously described by Nogueira-Campos et al (2012). These parameters were related either to the cell body or to the dendritic tree.

Cell body parameters included cell body area (corresponding to the flat 2D surface occupied by the neuronal soma) and *form factor* (that indicates how spherical the cell body is). This value varies between 0-1 and the closer to 1 (the maximum value), more spherical is the cell body.

Regarding to dendritic arborization, the morphological parameters studied were: (1) dendritic length, corresponding to the sum of the length of all segments of the dendritic tree; (2) number of nodes, corresponding to a total number of points that the dendritic tree originates two or more dendritic segments; (3) number of dendritic segments, corresponding to a total number of dendritic ramification between two nodes, or between a node and the cell body, or to a terminal branch; (4) fractal dimension; and (5) dendritic field area (after Wassle & Boycott, 1991). The fractal dimension gives a quantitative estimate of the complexity of the dendritic arborization, describing the way the dendritic ramification fills the area that comprised the dendritic field. The (5) area of the dendritic field was accessed by convex hull analysis, which measures the size of the dendritic field by interpreting a branched structure as a solid object in a given volume or area. The (6) dendritic orientation of each neuron was defined using wedge analysis of dendritic length (dl). A Cartesian coordinate reference frame dividing eight equiangular wedges was centered at the cell body, with its principal vertical axis oriented toward the pia mater. The total length of the dendritic segments contained in each wedge (dln) was measured to calculate a verticality index (*vi*) in which the sum of the dls in the four wedges close to the vertical axis was divided by the sum of dl measured in all eight wedges (total dl). The *vi* allowed the classification of type 1 neurons as horizontal- (0 < *vi* < 0.32), multipolar- (0.33 < *vi* < 0.65) or vertical-oriented (0.66 < *vi* < 1).

### Data analysis

The NN of three sequential slices of each animal was digitally reconstructed and their morphological parameters were compared both between the different cortical areas (M1 and S1) and the different laminar compartments of these two areas. The analysis was preceded by a Shapiro Wilk’s W normality test, and was found that the samples did not have a normal distribution. Therefore, the Kolmogorov-Smirnov two-sample test was performed to access statistical analysis. The level of significance was set at 0.05 in all analyses. Data from the different morphological parameters from each reconstructed neuron presenting a putative complete dendritic arborization in all histological sections analyzed were tabulated in a spreadsheet and transferred to a Systat 8 file (Systat Software, Inc., Chicago, IL, USA) for statistical analysis.

## 3. Results

In this study, we analyzed the distribution and morphology of the cell body and dendritic arborization of the NN in the representation of the forepaw in the S1 and M1 cortices of rats, in order to describe and compare them between these cortical areas and their compartments.

### Pattern of NADPHd-reactive neuropil staining

In the S1, the forepaw representation (region S1FL of Zilles, 1985) has an anteroposterior extension of about 2.5 mm. However, only in the most anterior 1.0 mm does forepaw border the primary motor cortex (the M1, or Fr1 area - Zilles, 1985). This was the region of interest in the present study because the M1 region contains the motor representation of the forepaw (Tandon et al., 2007). Thus, the sensory and motor representations of the forepaw neighbor each other in the same coronal section, being thus available for direct comparison after incubation in the same solution to reveal NADPHd reactivity. In this region, the S1 was easily identified due to the intense NADPHd reactivity of cortical layer IV, which stood out when compared with the UL and LL (Figure 1) and with the neighboring area M1 that did not present layer IV. Differences in the staining intensity of the diffuse NADPHd-reactive neuropil in the S1 additionally revealed the cortical layer borders. Typically, layer I corresponded to a thin densely stained layer close to the cortical surface followed by the less reactive layers II and III. The border between layers II and III could not be reliably identified by means of NADPHd histochemistry. Below the strongly reactive layer IV, there was the less reactive layer V. Layer VI, although clearly more intensely stained than layer V, was not as reactive as layer IV (Figure 1c).

In a typical coronal section, as one progresses from S1 towards the midline, the transition to area M1 is characterized by the rather abrupt termination of layer IV of the S1 (Figure 1c), giving way to a more homogenous pattern of neuropil staining, in which UL (layers I, II and III), plus LL (layers V and VI), display a similar pattern of staining as that of the S1. In the M1, NADPHd-reactive neuropil in layers I, II and III produce a uniform and moderate staining, more intense than the one found in the subjacent layer V. In addition, as in the S1, layer VI in the M1 is more intensely stained than layer V. However, in the S1, layers V and VI together are slightly more intensely reactive than their counterparts in the M1 (Figure 1c).

### Distribution and density of NN in S1 and M1

In order to quantify the spatial distribution of NN in the different cortical compartments of the S1 and M1, neuronal density was calculated in each hemisphere after adding up the number of cell bodies and the area size of individual compartments in all sections of the S1 and M1. Since simple cell counting may result in biased estimations of the number of cellular profiles (Gundersen et al., 1988), appropriate stereological corrections were adopted (see Materials and Methods section). In the hemisphere studied (the left), when compartment areas were added together from all the 150 μm-thick sections of the S1 or M1, the total area of the representation of the forepaw in the S1 ranged from 7.47 mm^2^ and from 7.26 mm^2^ for M1. The S1 and M1 comprised 60% LL and, correspondingly, these layers contained from 50 to 60% of all S1 or M1 NN. However, the highest nitrergic neuronal density values were found in layer VI in the S1 (about 165 neurons/mm^3^) and in the UL of the M1 (149 neurons/mm^3^). Overall, the neuronal density of the S1 (130 neurons/mm^3^) was greater than that of the M1 (119 neurons/mm^3^). The layer V had the lowest density values in all cases. Table 1 shows the results of this analysis.

**Table 1.** Distribution of nitrergic neurons cell bodies in the different cortical compartments.

| Cortical compartments* | Number of cells | | Compartment area (mm<sup>2</sup>) | | Cell body diameter (variance) in μm | | Neuronal density (cells/mm<sup>3</sup>)** | |
| --- | --- | --- | --- | --- | --- | --- | --- | --- |
|  | S1 | M1 | S1 | M1 | S1 | M1 | S1 | M1 |
| Total area | 158 | 141 | 7.47 | 7.26 | 12.26 (2.8) | 12.64 (2.6) | 130 | 119 |
| UL (I-III layers) | 45 | 71 | 1.77 | 2.93 | 12.65 (2.6) | 12.7 (2.0) | 156 | 149 |
| GL (IV layer) | 21 | - | 1.03 | - | 13.14 (1.3) | - | 125 | - |
| LL (V-VI layers) | 92 | 70 | 4.68 | 4.33 | 11.88 (2.8) | 12.58 (3.3) | 121 | 99 |
| Layer V | 34 | 42 | 2.5 | 2.6 | 12.40 (2.8) | 12.72 (3.3) | 84 | 99 |
| Layer VI | 58 | 28 | 2.17 | 1.73 | 11.57 (2.6) | 12.36 (3.2) | 165 | 100 |

\* Cortical compartments: UL, upper layers; GL, granular layer; LL, lower layers; \*\* Corresponding values were calculated after stereological corrections described in section "Stereological Evaluation of Cell Density" (see Materials and Methods section).

### Morphology of NN

We reconstructed all NADPHd neurons within the defined areas of the S1 and M1 in three sequential sections for two rats (Figure 2a and b). The NN located at the edges of the areas that showed more than half of the body within the sector were analyzed in the study, and the ramification of the NN were analyzed irrespective of whether parts of the neurites were located outside the boundaries of the sector where the neuronal body was. For the analysis, neurons were divided into eleven sectors, six in the S1 and five in the M1. They were: (a) the entire cortex, including all layers; (b) the upper layers [UL, layers I-III]; (c) granular layer [GL, layers IV, exclusive for the S1]; (d) layer V; (e) layer VI; (f) lower layers [LL, layer V and layer VI].

**FIGURE 2.**
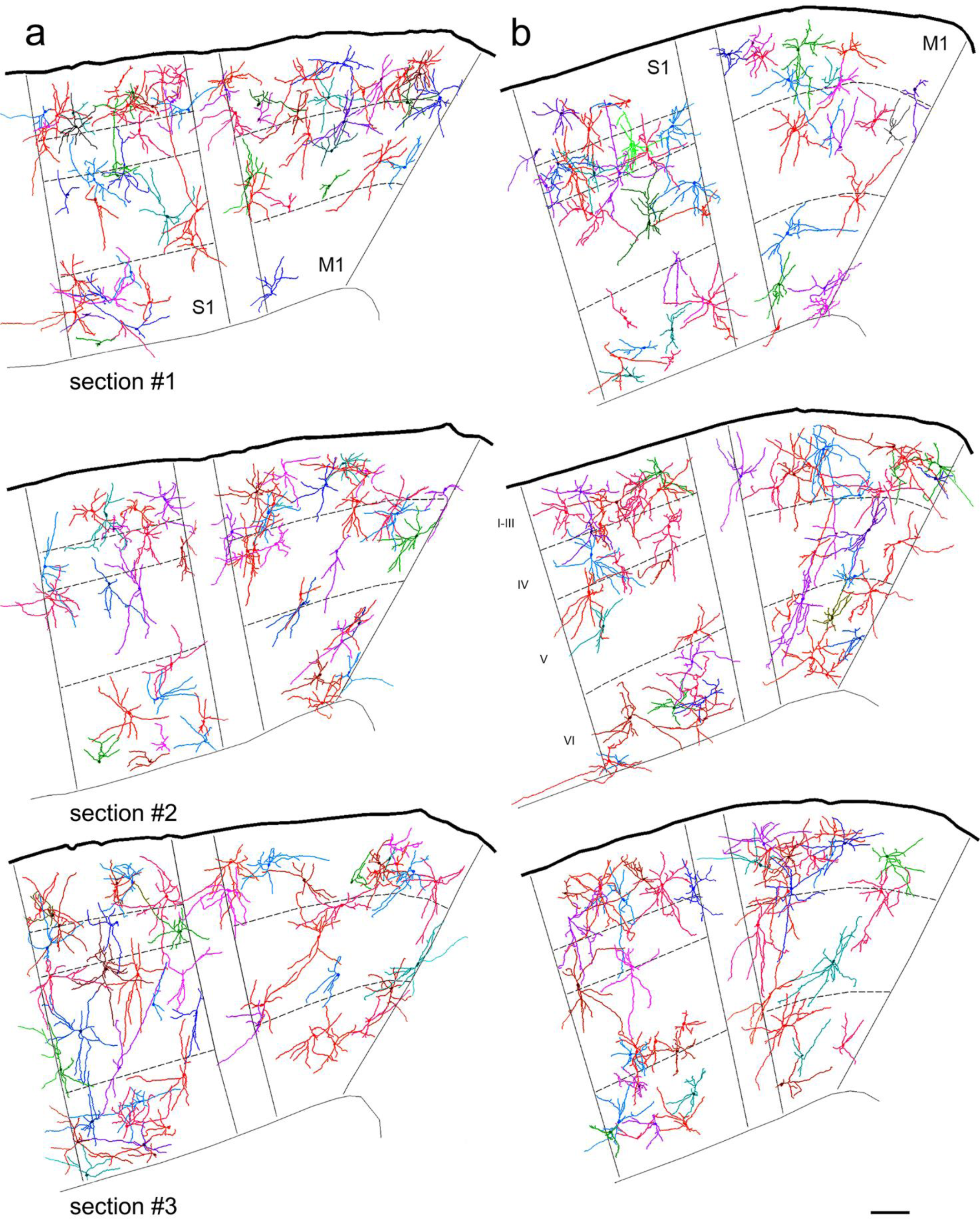
3D-reconstruction of all nitrergic neurons found in the representation of the forepaw of the rat in different cortical compartments in S1 and M1. All nitrergic neurons (NN) are oriented in the figure as if the pia mater were parallel to the top in the three sections of the case one (a) and case two (b). NN are more densely packed in the upper layers (I-III) and more sparsely distributed in layer V. Cell body distribution in S1 is different from that in M1. In S1, cells in lower layers are more densely packed in layer VI than in layer V. In M1, most cells are in the upper layers, while in S1, cells are both in upper layers and layer VI. Scale bar: 100 μm.

We were able to collect all the histological sections along the S1 and M1 without significant tears or tissue loss. The left hemisphere of these sections were thus used to perform a delimitation of rat S1 and M1, including 3D complete reconstructions of all NN located in all these areas in the forepaw representation, as illustrated in Figure 2 and 7. However, because some dendrites were cut during tissue sectioning, quantitative parameters were extracted from 209 NN located at mid-section depth and selected based on apparent completeness of all their dendrites.

**FIGURE 3.**
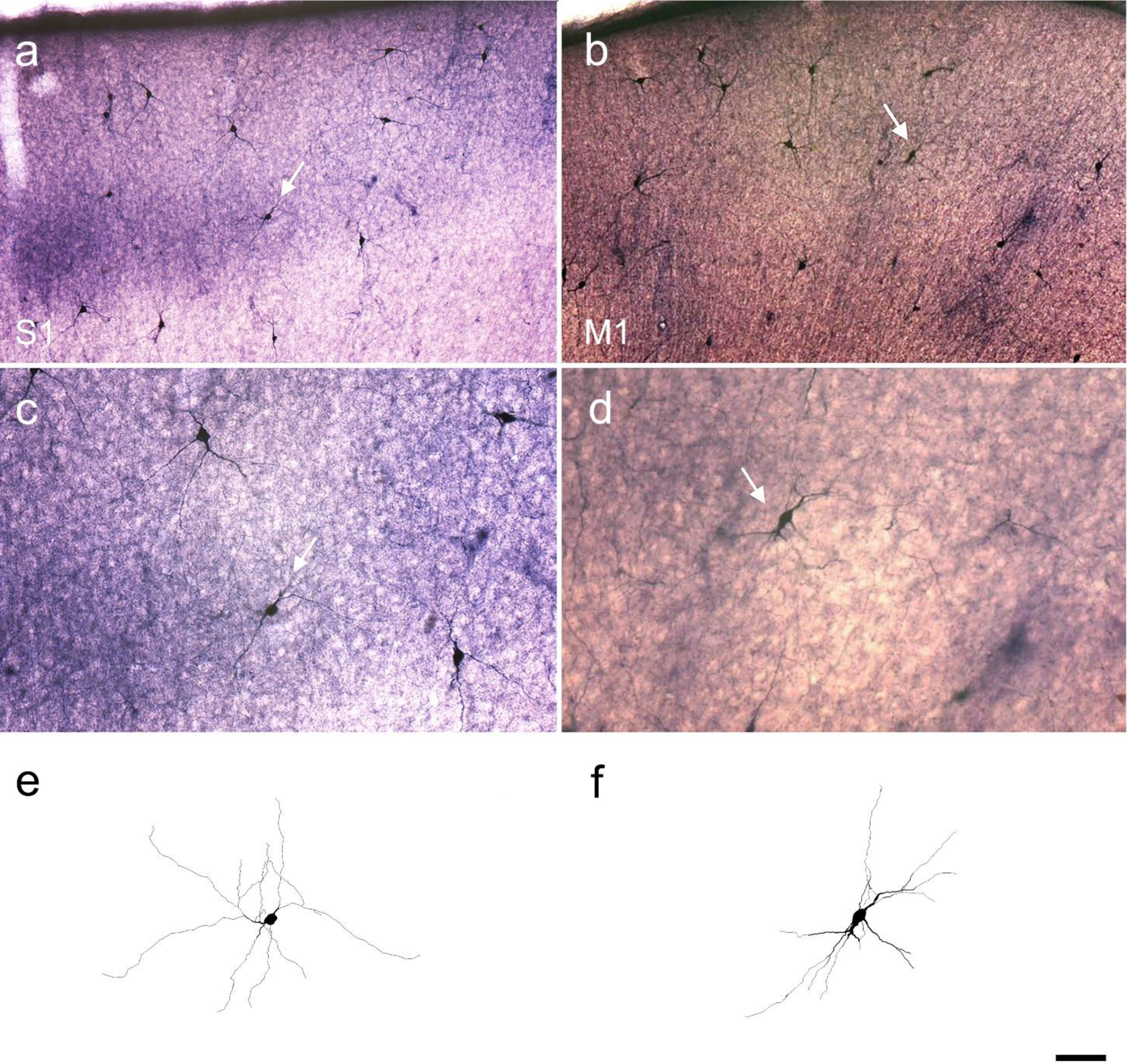
Photomicrographs and 3D-reconstructed NADPHd labeled neurons throughout areas S1 and M1. (a) A nitrergic neuron (NN) in layer IV in S1, and in the upper layers (UL) in M1 (b). (c) Neuron shown in (a), at high magnification. (d) Neuron shown in (b), at higher magnification. (e) 3D-reconstruction of the neuron highlighted in S1 (c), (f) 3D-reconstruction of the neuron highlighted in M1 (d). White arrows point to NN in the cortex. Scale bar: (a) and (b): 100 μm. (c), (d), (e) and (f): 50 μm.

**FIGURE 4.**
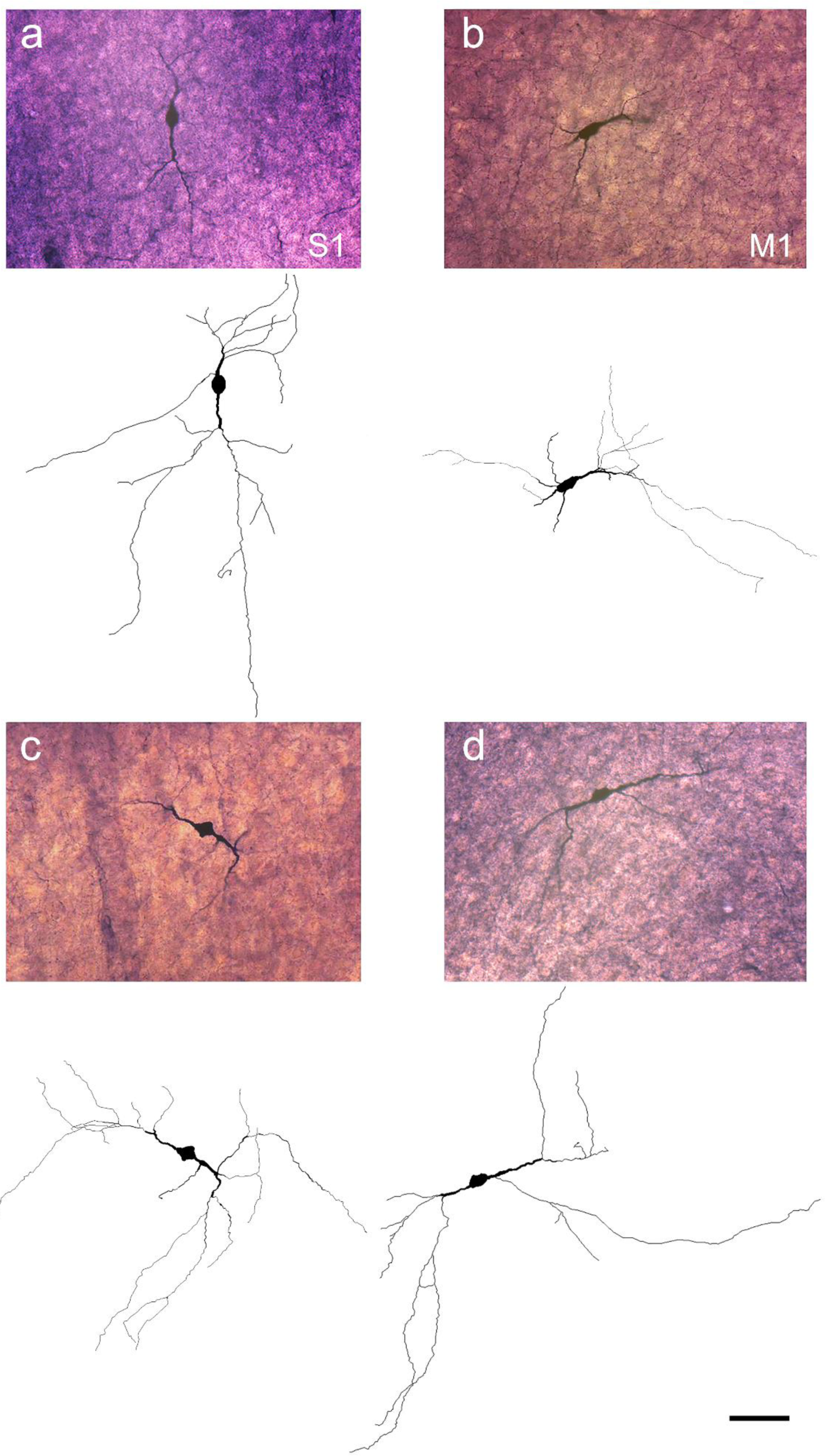
Photomicrographs and 3D-reconstructions of NADPHd labeled neuron in different cortical layers in S1 and M1. (a) Photomicrograph and reconstruction of a nitrergic neuron (NN) located in upper layers in S1; (b) NN located in upper layers in M1; (c) NN located in lower layers in S1; and (d) NN located in lower layers in M1. Scale bar: 50 μm.

**FIGURE 5.**
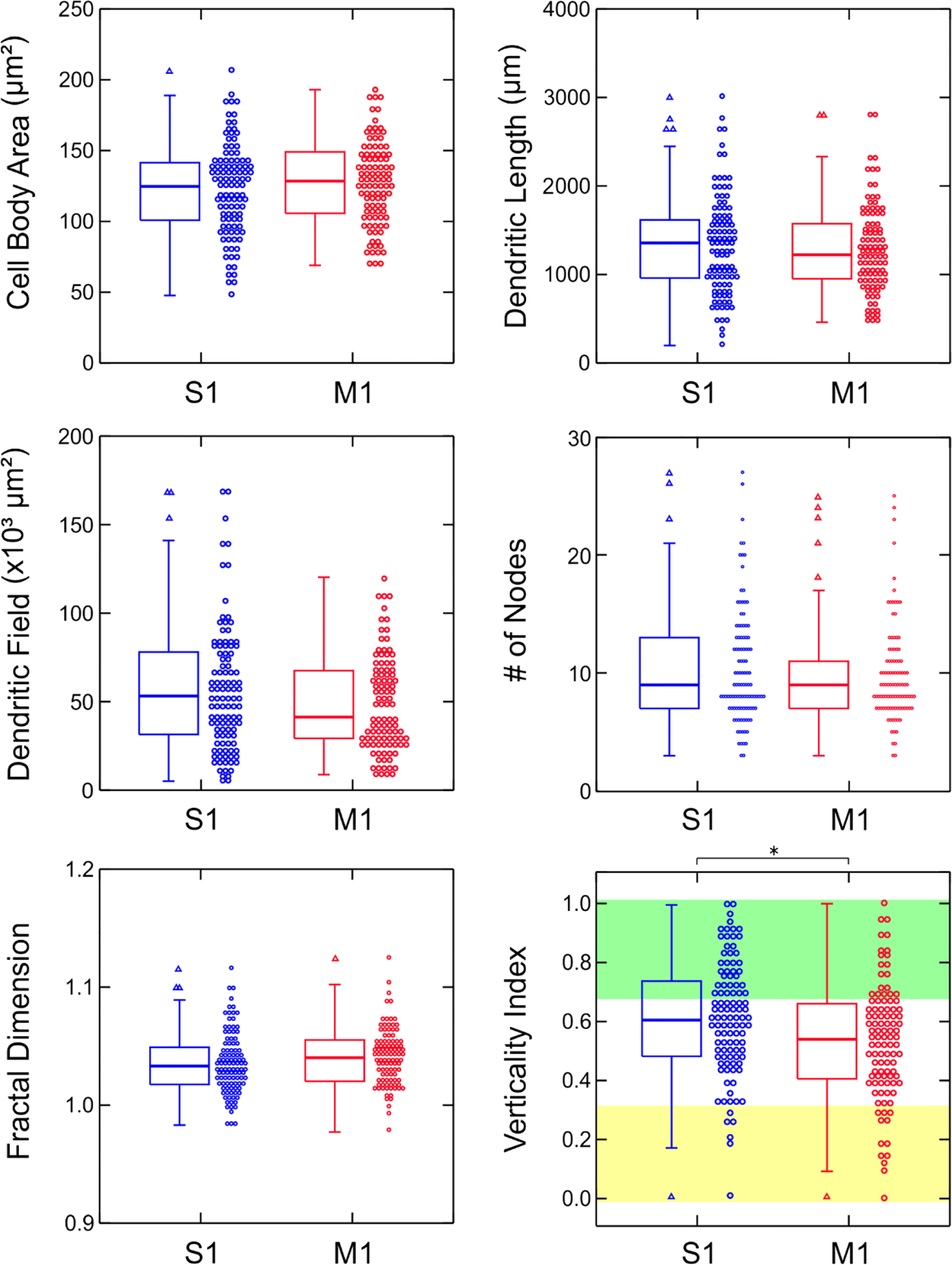
Comparison of morphological parameters of nitrergic neurons of S1 and M1. Median (heavy line), first and 3rd quartile (box), maximum and minimum (vertical lines), and individual values (circles) of cell body and dendritic parameters. In the verticality index graph, the yellow band corresponds to horizontal neurons (vi= 0 to 0.32), the white band (vi=0.33 to 0.65) to multipolar neurons, and the green band to vertical-oriented type 1 neurons (vi=0.66 to 1). S1 in blue, and M1 in red. Asterisk (*) represents a significant difference of p < 0.05; triangles are outliers.

**FIGURE 6.**
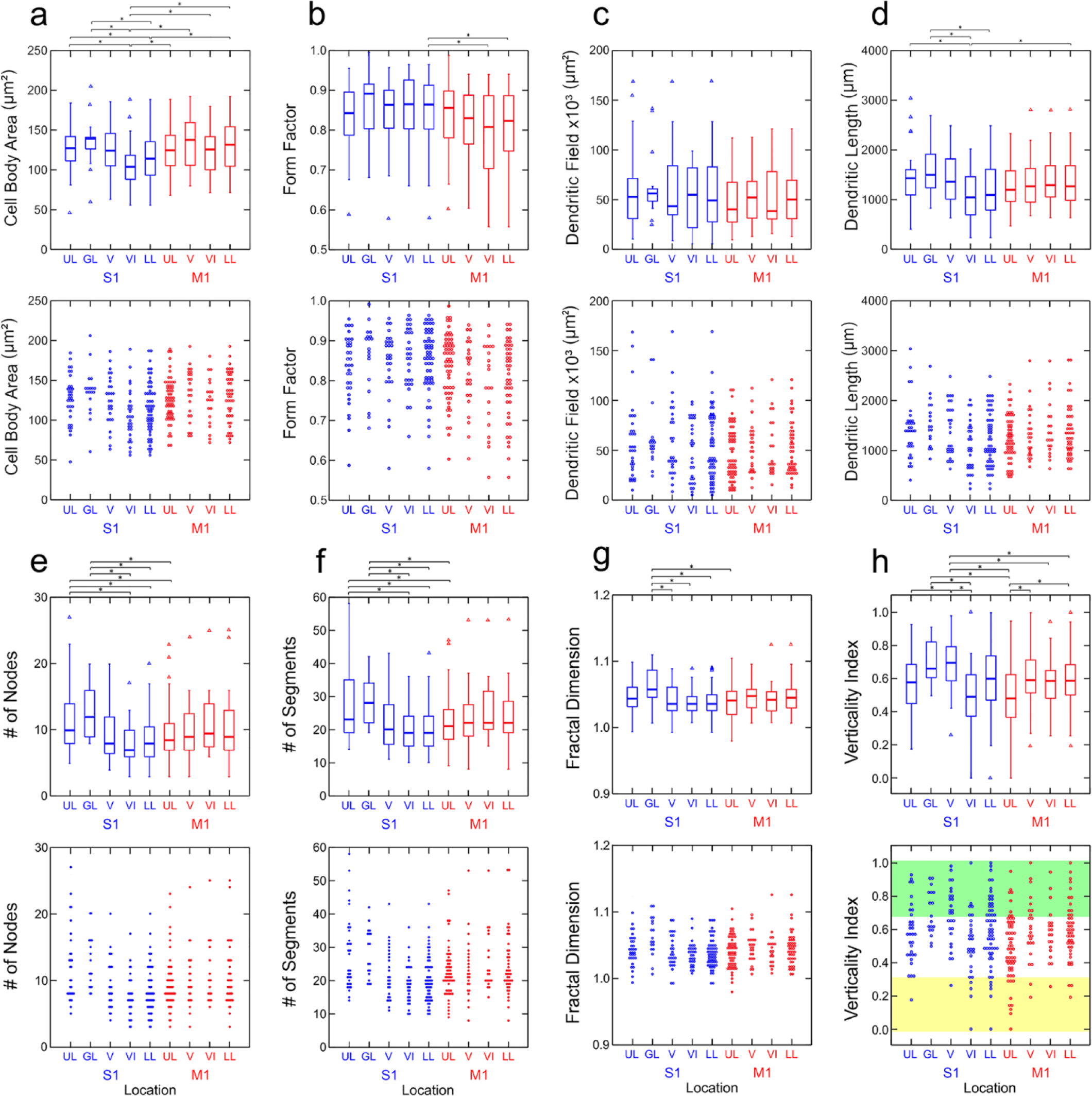
Comparison of morphological parameters of nitrergic neurons (NN) in different cortical compartments of S1 and M1. Median (heavy line), first and 3rd quartile (box), maximum and minimum (vertical lines) are illustrated in the top and individual values (circles) are shown in the bottom. Values for cell body area (a), form factor (b), dendritic field (c), dendritic length (d) number of nodes (e), number of segments (f), fractal dimension (g) and verticality index (h) of NN are presented for upper layer (UL), granular layer (GL), layer V (V), layer VI (VI), and lower layers (LL). S1 in blue, and M1 in red. In the verticality index graph (H), the yellow band corresponds to horizontal neurons (vi= 0 to 0.32), the white band (vi=0.33 to 0.65) to multipolar neurons, and the green band to vertical-oriented type 1 neurons (vi=0.66 to 1). Asterisk (*) represents a significant difference of p < 0.05; triangles are outliers.

**FIGURE 7.**
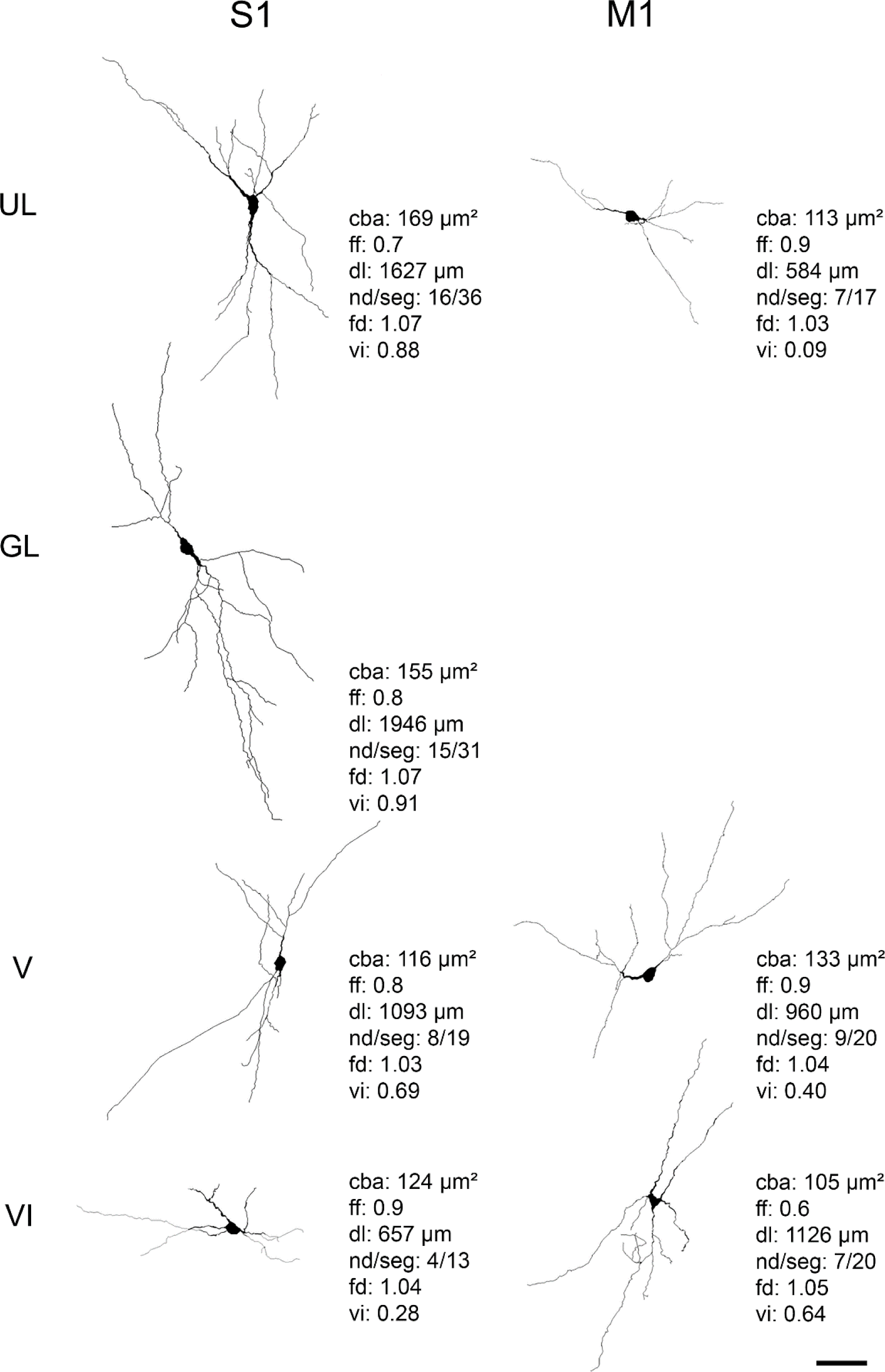
Examples of 3D-reconstruction of typical nitrergic neurons from different cortical compartments. All nitrergic neurons (NN) are oriented as if the pia mater were parallel to the top of the figure. NN in the left were respectively located in S1 upper-layers (UL), granular layer (GL) and lower layers V and VI. NN in the right were respectively located in M1 upper-layers (UL) and lower layers V and VI. The NN in the S1 display more vertically-oriented dendritic trees, while M1 NN tend to be horizontally-oriented or multipolar. In S1: UL neurons have larger cell body area than lower layers (LL) neurons. GL neurons have greater dendritic length than LL neurons. In M1: LL neurons have greater vertically-oriented dendritic trees than UL neurons. Values of four relevant morphological parameters are depicted for each neuron illustrated: cba, cell body area; ff, form factor; dl, total dendritic length; nd, nodes; seg, segments; fd, fractal dimension; vi, verticality index. Scale bar: 50 μm.

Figure 3 shows two similar type I neurons from the S1 and M1 with different cell bodies located in a central position and with different dendritic arborization. The neuron from the S1 is located in layer IV while that from the M1 is located in UL.

Figure 4 shows four different type I neurons from the UL and LL of the S1 and M1. These neurons are differentially oriented in relation to the pial surface, have different cell body areas, are asymmetrically located within the dendritic arborization, and have different dendritic field areas. The first two neurons from the S1 (a) and the M1 (b) are located in UL, while the remaining ones are from the LL.

### Quantitative analysis of the NN morphological parameters

Morphological parameters of the cell body (cell body area and *form factor*) and dendritic branching (dendritic length, dendritic field, number of nodes and segments, fractal dimension and verticality index) of the NN of the S1 were compared with those of the M1 and their different compartments. The comparisons are expressed as median, 1^st^ and 3^rd^ quartile, maximum and minimum (left) as well as individual values (right).

The Figure 5 shows that the NADPHd positive neurons found in the S1 have a higher verticality index than those of the M1 (p<0.05). There was no statistical difference for cell body area, dendritic length, dendritic field, number of nodes, or fractal dimension for cells from the forepaw representation of the S1 and M1.

Figure 6 shows a quantitative evaluation of the statistical differences in the morphological parameters of the different S1 and M1 cortical compartments (p<0.05). Figure 6a shows the statistical differences in cell body area in the S1 (blue) and M1 (red) cortical compartments. The sample was divided into five S1 compartments, namely: upper layer (UL), granular layer (GL), layer V (V), layer VI (VI) and lower layer (LL): while the M1 was divided into four compartments: upper layer (UL), layer V (V), layer VI (VI) and lower layer (LL). With respect to cell body, the NADPHd neurons in the UL of the S1 have larger areas than the neurons in the LL of the S1. This difference is particularly significant when comparing the UL with layer VI. In addition, neurons in layer VI of the S1 have a smaller cell body area than those of the UL and GL, or the LL of the M1. The neurons of the LL of the M1 have a larger cell body area than the neurons in the same compartment in the S1.

Figure 6b shows the statistical differences in the form factor of the cell body neurons found in the different S1 and M1 cortical compartments. The neurons found in the LL of the S1 are more spherical than those from layer VI or the LL of the M1

Regarding dendritic length (Figure 6d), neurons in the UL of the S1 have a greater total dendritic length than the neurons of layer VI from the same cortical area. The neurons of the GL have a greater dendritic length that those from layer VI or from LL of the S1. In turn, neurons in layer VI of the S1 have a smaller total dendritic length than those of the LL of the M1. Figure 6c shows no statistical differences in the dendritic field area of the neurons found in the different S1 and M1 cortical compartments (p< 0.05).

Figure 6 also shows the statistical differences in the number of nodes (Figure 6e) and the number of segments (Figure 6f) of the neurons found in the S1 and M1 cortical compartments (p<0.05). The neurons in the UL of the S1 have more nodes than those of the neurons in layer VI or in the LL of the same cortical area. The neurons in the GL have more nodes than those of the neurons in layer VI or in the LL of the S1. In turn, the neurons in the UL of the S1 have more nodes those of the neurons from the same layer of the M1. In addition, the neurons in the GL of the S1 have more nodes than those of the neurons in the UL of the M1. Regarding the number of segments, the neurons in the UL of the S1 have a greater number of segments than those of the neurons in layer VI or in the LL of the same cortical area. The neurons in the GL have a greater number of segments than those of the neurons in layer VI or in the LL of the S1. In turn, the neurons in the UL of the S1 have a greater number of segments than those of the M1. The neurons of the GL of the S1 have a greater number of segments than those in the UL of the M1.

Figure 6 also shows the statistical differences in fractal dimension (Figure 6g) and the verticality index (Figure 6h) of the neurons found in the S1 and M1 cortical compartments (p<0.05). The neurons in the GL of the S1 have a greater fractal dimension than those of the neurons in layers V, VI or the LL of the S1. They also have a greater fractal dimension than the neurons in the UL of M1. With respect to the verticality index, the neurons in layer V of the S1 have a greater verticality index than the neurons in the UL and in layer VI. The neurons in the GL have a greater verticality index than those in layer VI of the S1. In the M1, the neurons in the UL have a lower verticality index than those in layers V or LL of this area. In addition, neurons in the GL of the S1 have a greater verticality index than those in the UL of the M1. Furthermore, the neurons in layer V of the S1 also have a greater verticality index than the neurons in the UL, layer VI or LL of the M1.

The Table 2 describes the distribution of the NN according to their dendritic orientation and shows that the majority of the NN are multipolar and vertical (33% and 10%, respectively) and the minority (6%) were horizontal. In S1, 58% of the NN were multipolar and 36% were vertical, and a minority were horizontal (6%). In M1, the majority of the NN were multipolar (69%), the vertical neurons (20%) and horizontal NN (12%). Regarding S1 and M1 compartments, the multipolar NN of the S1 are more common in the UL (70%), horizontal neurons in layer VI (14%) and vertical NN in layer V (56%). In M1, multipolar NN are mostly in layer VI (75%) followed by upper layers with a percentage of 71%, vertical NN of the M1 are more common in layer V and horizontal neurons in UL.

**Table 2.** Distribution of nitrergic neurons according to their dendritic orientation

|  | Vertical |  |  | Multipolar |  |  | Horizontal |  |  |
| --- | --- | --- | --- | --- | --- | --- | --- | --- | --- |
|  | S1 | M1 | Total | S1 | M1 | Total | S1 | M1 | Total |
| **Total area** | 39 (36%) | 20 (20%) | 20 (10%) | 62 (58%) | 70 (69%) | 70 (33%) | 6 (6%) | 12 (12%) | 12 (6%) |
| **UL (I-III layers)** | 9 (27%) | 8 (14%) | 8 (9%) | 23 (70%) | 41 (71%) | 41 (45%) | 1 (3%) | 9 (16%) | 9 (10%) |
| **GL (IV layer)** | 9 (50%) | - | - | 9 (50%) | - | - | 0 (0%) | - | - |
| **LL (V-VI layers)** | 21 (38%) | 12 (27%) | 12 (12%) | 30 (54%) | 29 (66%) | 29 (29%) | 5 (9%) | 3 (7%) | 3 (3%) |
| **Layer V** | 15 (56%) | 8 (33%) | 8 (16%) | 11 (41%) | 14 (58%) | 14 (27%) | 1 (4%) | 2 (8%) | 2 (4%) |
| **Layer VI** | 6 (21%) | 4 (20%) | 4 (8%) | 19 (66%) | 15 (75%) | 15 (31%) | 4 (14%) | 1 (5%) | 1 (2%) |

\* Number of nitroergic neurons (NN) per compartment and percentage of the total number of NN reconstructed in the sample = 209
NN; Total: sum of NN of the S1 and M1.

## 4. Discussion

We described and compared the distribution and morphological arrangement of type 1 NN neurons in the forepaw representation in the S1 and M1 of the adult rat. These neurons are well known to release NO, a small gaseous molecule involved with various fundamental physiological and pathological roles in the central nervous system (Bredt & Snyder, 1992; Dawson et al., 1994). S1 and M1 have distinct functionalities and are adjacent areas in the cortical forepaw representation of the rat. They are composed of different cortical compartments representing different laminar arrangement topologically organized (Donoghue et al., 1990; Donoghue & Wise, 1982; Lübke & Feldmeyer, 2007; Tandon et al., 2007; Wester & Contreras, 2012), characteristics that were explored in the present study.

### Distribution and density of NN in S1 and M1

We found differences in the distribution and density of the NN across the forepaw representation in the S1 and M1. The neuronal density of the S1 were greater than that of the M1. Similar results were found in the distribution of these neurons in the S1 and M1 whisker representation (Afarinesh & Behzadi, 2018) and in these same cortical areas including other cortical representations (Huh et al., 1998).

We demonstrated the highest nitrergic neuronal density values in layer VI in the S1, according to previous study in the whisker representation in the S1 (Afarinesh & Behzadi, 2018). When compare UL (including layers I-III), GL and LL (including layers V and VI), the highest nitrergic neuronal density values were in UL, consistent with Nogueira-Campos et al (2012) when observed the entire S1 cortex, including other parts of this cortical area like anterior snout, whiskers, forepaw, hindpaw and trunk. Other studies on distribution and density of NN mainly performed on tangential section along layer IV of the posterior medial barrel subfield (PMBSF, a region where whiskers are represented and the barrels are sharply-defined) to estimate the laminar distribution of the NN in S1. Such studies reveled higher distribution in the S1 supragranular layers and layer VI, similar results with those of this study (Franca et al., 2000; Freire et al., 2004).

Regarding the distribution and density of NNs in forepaw representation in the M1, we found the highest neuronal density in UL followed by layer VI. Other studies found the most NN located in layers V (in the M1 whisker representation, Afarinesh & Behzadi, 2018) and in layer VI of the M1 (Vlasenko et al., 2008), but the last study does not explain which parts of this cortical area were contained in the sections selected. These distinct results may be associated with different cortical representations of each of these studies. Future researches are necessary to better understand the NO role in the M1 and the cortical physiology.

### Nitrergic neuronal morphology in S1 and M1

#### a) Cell Body

The morphology of the NN were evaluated in relation to their cell body area and shape (*form factor*). When we compared the cell body area and *form factor* of the entire two different cortical areas (S1 and M1), we did not find any statistical differences. Afarinesh & Behzadi, 2018 reported different results in the whisker representation: the mean soma diameter of NN in the M1 was less than NN in S1. We revealed that the neurons of the LL of the M1 have a larger cell body area (but less spherical) than the neurons in the same compartment in the S1 (Figure 7). This may be associated with the relevant role of the lower layers of M1, which participate in the generation of motor commands and coordinated movements (Petersen, 2019), and the perimeter of the cell body can increase at the expense of its sphericity to increase the supply of NO. Additionally, the cell body morphometry may vary across different cortical representations in S1 and M1.

In the S1 representation of the forepaw, the NN found in the UL of the S1 had a larger cell body area than those in the LL. Particular significance were found when compare UL and GL with layer VI: NN in layer VI have smaller cell body area than those of this other layers (Figure 7). Nogueira-Campos et al (2012) showed that NN in the GL have larger cell body area than those in other layers along S1, similar with this study. These results are strengthens with data in the literature that observed in primary sensory areas such as S1, a high metabolic activity in GL that can be identified by intense histochemical reactivity (Wallace, 1987; Wong-Riley & Welt, 1980). This high concentration of metabolic enzymes (such as succinate dehydrogenase and cytochrome oxidase) in GL correlates with an intense reactivity to NADPHd and NOS (Aoki et al., 1993; Wong-Riley et al., 1998), proposing that NO synthase expression is coupled to neural activity (Garthwaite, 1995). The role of NO release in the neuropil need to be further investigated in other studies.

#### b) Dendritic arborization and orientation

We analyze the dendritic arborization of NN in the representation of the forepaw in the entire two different cortices (S1 and M1) and did not find any statistical differences. In the representation of the whiskers, Afarinesh & Behzadi (2018) found significant distinctions between S1 and M1 dendritic arborization: NN in the M1 had fewer 3^rd^ order dendritic processes, nodes and segments. In the present study, we found difference between the S1 and the M1 cortical compartments: NN in the UL of the S1 have more nodes and segments than those of the neurons from the same compartment of the M1. Neurons in the GL of the S1 have greater fractal dimension, more nodes and segments than those of the neurons in the UL of the M1 (Figure 7).

In S1, when the morphological parameters related to the dendritic tree were analyzed along the cortical compartments, we reveled differences in the dendritic length, number of nodes and segments, and fractal dimension of the NN (Figure 7). Those located in GL of S1 have greater dendritic length, number of nodes, number of segments and fractal dimension than those in the LL and layer VI of the same area. The NN in the UL of the S1 had greater dendrite length, number of nodes and segments than NN in the layer VI, and have greater number of nodes and segments than NN in the LL. Additionally, differences in the dendritic arborization did not present across compartments of M1.

The analysis of the dendritic orientation of the NN in the representation area of the forepaw in the entire S1 and M1 showed that most NN are multipolar and vertically-oriented and the NN located in the S1 have a higher vertical index than those of the M1. With respect to the NN in the S1 and M1 cortical compartments, the NN in the GL of the S1 presented a greater verticality index than those of the UL of the M1. The NN in the layer V of the S1 have the greatest verticality index than UL, layer VI and LL of the M1 (Figure 7).

We comparing the orientation of the dendritic tree of the NN of the forepaw representation in the S1. We demonstrated that a minority of NN is horizontal (with a percentage of 6%), according to Nogueira-Campos et al (2012) that found 7-26% horizontal NN when the whole S1 was analyzed.

We also demonstrated that the majority of the NN in the S1 are multipolar (a percentage of 58%) and vertically-oriented (a percentage of 36%), but Nogueira-Campos et al (2012) found in the entire S1 50% of NN with dendritic trees vertically oriented and 24-29% multipolar (Table 2). Nogueira-Campos et al (2012) revealed in the entire S1 that multipolar NN were more common in the UL, according to this study that found a percentage of 70% of multipolar NN located in this same compartment. Additionally, we found a predominance of vertical NN in the UL compartment of the S1 than horizontal neurons, in disagreement with what was report for entire S1 (Nogueira-Campos et al, 2012).

We demonstrated that NN in layer V have greater verticality index than those in the UL and in layer VI of the S1. Also revealed that the NN in the GL have greater verticality index than those in layer VI of the S1, but not than LL (with layer V e VI together), Figure 7. Nogueira-Campos et al., (2012) found that the verticality rates were significantly lower in the UL than GL and LL of the S1 but did not investigated de LL separately. In addition, we also found that the M1 neurons in the UL have a lower verticality index than those of layer V and the LL of this same area (Figure 7).

We propose that differences in the cell body and dendritic parameters of the NN may differ across the different cortical representation and in different cortical layers in the forepaw representation in the S1 and M1. The larger cell body as well as the greater length, fractal dimension, nodes and segments of the dendritic tree could be associated with higher concentration of the enzymes that synthase NO in association with the local neural activity in the layer GL that is exclusive to the S1. As NN are involved in neurovascular coupling (Cauli, 2010; Drake & Iadecola, 2007), while the S1 non-vertical NN in UL could participate maintain the metabolic demands necessary for horizontal integration and the NN in GL and layer V can simultaneously maintain the vertical activity with those significant projections in the cortex of the rat (Schubert et al., 2007).

### Technical considerations

We used the same perfusion and fixation procedures in both the animals, but variations in the fixation intensity are usually very difficult to avoid and may have occurred in our material (Matsumoto et al., 1993). Romanelli et al (2007) have reported an excess of nitrergic expression that can be reversible regulated in the S1 in rat brains after being exposed to a new location. Moreover, the morphological complexity of NN may be regulated by many factors related with the phenotypic variation that can be present in different animals, same determinants are: physical exercise (Torres et al., 2006), light exposure (Chen et al., 2013), and emotional stress (Beijamini & Guimarães, 2006). In addition, there are subpopulations of NN, such as NN co-localized with neuropeptide Y (Estrada, 1998), that in this study are being analyzed together, without comparing the morphology of each of them separately.

Moreover, Hilbig and Punkt (1997) showed day-time-dependent changes of NADPHd activity in the neuropil of the visual cortex. We therefore, performed the perfusion at the same time of day, between 7am and 8am, the period of maximum enzyme activity in the rat cortex.

## 5. Conclusion

The forepaw representation of the S1 have greater neuronal density than M1. On the other hand, NN did not differ their cell bodies and dendritic arborization morphology when these two areas were compared without considering their distinct laminar arrangements. However, the morphology of the cell body and dendritic tree of NN differs between different cortical compartments in the forepaw representation in the S1 and M1.

In the S1 and M1 forepaw representation of the rat, the NN correspond to a single neuronal population whose functionality is independent of the peculiarities of the different types of sensory and motor processing. However, the morphological differences found between cortical compartments of these areas, as well as the higher density of NN found in the S1 indicate that the release of NO varies between the S1, M1, and their compartments.

### In memoriam of João Guedes da Franca

This paper was done under the supervision of Professor João Guedes da Franca and his contribution to this paper was fundamental. He was a great young scientist who left us too soon. João Franca made important contributions to the anatomy of the cerebral cortex in several species. His great scientific expertise and rigorous application of anatomical and electrophysiological techniques will be always remembered.

## Authors contributions

All authors had full access to all the data in the study and assume responsibility for the integrity of the data and the accuracy of the data analysis. Study concept and design: MRC, JCH, AANC, JGF. Date acquisition: BPP, AAN, MRC. Analysis and interpretation of data: BPP, RG, JGF. Drafting of the manuscript: BPP, RG. Critical revision of the manuscript for important intellectual content: JGF, RG, JCH, MRC, AANC. Statistical analysis: BPP, RG. Obtaining funding: JGF, RG. Study supervision: JGF, RG, JCH.

## Data Availability Statement

Data have not been shared. Our web repository is under construction. Data is available upon request to the first author.

## Abbreviations

NN: nitrergic neurons
NO: nitric oxide
NADPHd: nicotinamide adenine dinucleotide phosphate diaphorase
S1: primary somatosensory cortex
M1: primary motor cortex
NOS: nitric oxide synthases
PB: phosphate buffer
UL: upper layers
GL: granular layer
LL: lower layers
dl: dendritic length
vi: verticality index
PMBSF: posteromedial barrel sub-field

